# Load-dependent and load-independent effects on longitudinal motor training in human continuous hand movements

**DOI:** 10.1101/2024.07.01.601475

**Authors:** Xiaolu Wang, Xuan Liang, Yixuan Ku, Yinwei Zhan, Rong Song

## Abstract

Motor learning involves complex interactions between the cognitive and sensorimotor systems, which is susceptible to different levels of task load. While the mechanism underlying load-dependent regulations in cognitive functions has been extensively investigated, their influence on downstream execution in motor skill learning remains less understood. The current study extends the understanding of how load levels affect motor learning by a longitudinal functional near-infrared spectroscopy (fNIRS) study in which 30 participants (15 females) engaged in extensive practice on a two-dimensional continuous hand tracking task with varying task difficulties. We propose the index of difficulty (ID) as a quantitative estimate of task difficulty, which is positively correlated with psychometric measure of subjective workload level. Results shows that as behavioral performance improved over time, participants adopted a direction-specific and load-independent (i.e., consistent across different load levels) control strategy, shifting from feedback-dominant to feedforward-dominant control in the vertical direction as training progressed. Crucially, we provide robust evidence of the learning-induced alteration in load-dependent cortical activation patterns, suggesting that effective motor skill learning may lead to shift towards an inverted-U relationship between activation and load level in the pre-motor and supplementary motor areas. In addition, brain-behavior relationship in the frontoparietal network was strengthened after training. Taken together, our findings provide new insights into the learning-induced plasticity in brain and behavior associated with load-dependent and load-independent contributions to motor skill learning.

## Introduction

Humans’ ability to acquire and consolidate motor skills is prone to subjective load level. It has been suggested that optimal load level, regulated by task difficulty, could facilitate learning in both lab-based (e.g., force control, reaching and tracking movements) and real-world (e.g., driving, dance and sports) scenarios (Akizuki and Ohashi, 2015; Liao et al., 2018; Schaefer et al., 2021). This raises questions about how to determine the load level that maximizes learning outcomes, or in other words, how to characterize the non-monotonic (inverted-U) shaped relationship between load level and behavioral/neural metrics. Understanding the neural substrate of the load-dependent learning effects is therefore crucial for answering this question, which is a key goal in domains such as biomedical engineering, cognitive and behavioral neuroscience.

Existing evidence from both single neuron recordings and functional neuroimaging studies has revealed the relationship between neural response and load level, mediated by increasing, decreasing or inverted-U function in both cognitive (Rypma et al., 1999; Capa et al., 2008; Chen et al., 2008; Nagel et al., 2009; Heinzel et al., 2014; Van Snellenberg et al., 2015; Lamichhane et al., 2020; Iordan et al., 2020; Meidenbauer et al., 2021) and motor tasks (Akizuki and Ohashi, 2015; Bosse et al., 2015; Shuggi et al., 2017; Zheng et al., 2021). Basically, the patterns of load-dependent neural response are typically task- and region-specific, whereas the inverted-U shaped relationship is relatively less reported. This is possibly due to the fact that many studies have focused their investigation on a narrow range of load levels (three or fewer), which is insufficient to identify the ‘inflection point’ in the inverted-U curve where the neural response might peak. Expanding the range of load levels has revealed clearer evidence of the inverted-U pattern, particularly in studies that included more than five load levels (Van Snellenberg et al., 2015; Iordan et al., 2020; Lamichhane et al., 2020; Zheng et al., 2021).

However, the identification of the inverted-U patterns is nuanced, usually characterized by an increase at low loads and a subsequent decrease at high loads (Van Snellenberg et al., 2015; Lamichhane et al., 2020; Zheng et al., 2021). In a recent study on working memory training, Lamichhane et al. (2020) used model-based approach to test the linear versus non-linear (quadratic) effect of load levels on blood oxygenation level-dependent (BOLD) activity, providing more robust evidence of the inverted-U pattern in the lateral prefrontal cortex. By contrast, the load-dependent patterns seem to be different in a force control motor task, where the inverted-U profile is observed in low-level networks such sensorimotor and visual areas, rather than prefrontal cortex (Zheng et al., 2021). On the other hand, regarding the learning-induced effect, a longitudinal study on working memory reported right shift of activation peak after training, indicating learning-induced brain plasticity in cognitive learning (Iordan et al., 2020); yet, evidence on motor skill learning is still lacking.

Surprisingly, research regarding the load-dependent effect on human motor control almost exclusively focused on discrete movement (e.g., reaching, grasping, motor sequence, etc). While the well-established Fitts’s index of difficulty (ID) provides a means for quantifying load level in point-to-point goal-directed movement, it appears that discrete and continuous movements may be supported by different mechanisms (Huys et al., 2008; Schaal et al., 2004; Yang et al., 2021). Unlike discrete point-to-point movements, motor control of continuous movements more closely simulates motor skill acquisition in real-world scenarios. Although some behavioral and neurofunctional research has explored the effects of task difficulty on behavior and neural representation in continuous motor control by varying trajectory complexity, speed or feature dimension (Shull et al., 2017; Gaume et al., 2019; Haga et al., 2002), much remains to be elucidated about the basis of motor learning in continuous movement control.

The current study thus aims to provide a more comprehensive understanding of the load-dependent effect on learning-induced plasticity in brain and behavior in motor training of continuous hand movements. Here, we propose an experimental paradigm that quantitatively manipulating subjective load level in continuous target tracking tasks. Furthermore, we use psychometric measure to validate the effectiveness of the proposed approach in reflecting subjective workload level. In the longitudinal functional near-infrared spectroscopy (fNIRS) study, participants engaged in extensive practice on the continuous target tracking task with varying load levels over a 5-day training period. Through systematic analysis, our findings provide evidence of both load-dependent and load-independent accounts associated with effective motor skill learning.

## Materials and Methods

### Participants

Thirty healthy right-handed volunteers (15 females; age 22-26 years; mean age 24 years) participated in the study. All participants had normal or corrected-to-normal vision and reported no history of psychiatric and/or neurological disorder. Handedness was assessed by Edinburgh Handedness Inventory (Oldfield, 1971). The study was approved by Ethics Committee of Department of Psychology of Sun Yat-sen University and completed in accordance with the Declaration of Helsinki. All participants gave written informed consent and were compensated CNY 200, plus a bonus up to CNY 150 depending on performance.

### Experimental Design

#### Experimental setup

The experiment was conducted in a quiet and dimly lighted room to avoid distractions. Participants sat comfortably on a chair, with head restricted by a height-adjustable chin rest. Visual stimuli were presented on a 23.8-inch monitor (AOC, resolution: 1920 × 1080, refresh rate: 60 Hz) at viewing distance of 75 cm. No verbal instructions were given during the main tasks.

#### Procedure and task design

Each participant visited the laboratory for 5 consecutive days and finished 5 experimental sessions (1 session/day), including 2 evaluation sessions and 3 training sessions (Figure 1A). Prior to the main experiment, participants had a warm-up session that they could practice each experimental condition once. During evaluation sessions, fNIRS signals were recorded through out the whole experimental period. After each block, a 5-point (5-excessive, 4-high, 3-comfortable, 2-relaxed, 1-under utilized) instantaneous self-assessment (ISA) rating scale was presented to assess the subjective workload level(Tattersall and Foord, 1996). There were 60 s inter-block breaks after ISA ratings. During the training sessions, only the task blocks were retained, while all other components—including the 60-second inter-block breaks, fNIRS recordings, and ISA ratings—were removed. For incentivization, a ranking list organized by overall success rate was displayed following the completion of each session. Participants were informed that the top three performers in the experiment would receive an additional bonus after all participants had completed the experiment.

The experiment uses a within-subject design and follows block-design paradigm. During each session, participants performed 30 trials (6 difficulty levels × 5 repetitions), with 6 levels of task difficulty pseudorandomly presented in blocks. Each difficulty level condition consisted of 5 identical trials of 15 s, separated by 15 s fixation (Figure 1B).

Participants were instructed to control a circular cursor using a joystick (new T16000 FCS, Thrustmaster) with their right hand to track the movement of a black filled circle (the target) in two-dimensional pseudorandom motion as closely as possible.

The target movements in the horizontal (x-axis) and vertical (y-axis) directions are controlled separately by velocity vector,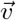. Specifically, the horizontal velocity is randomly selected from *v*_*x*_ *∈* {3, 6, 9, 12} (cm/s) every second, while the vertical velocity is determined by *v*_*y*_ = 4| sin (*t*)| · (15 *− v*_*x*_) (cm/s), which depends on time *t* (sampled at 60 Hz) and the horizontal velocity. Note that these parameters were set arbitrarily based on pre-experiment. Our main purpose is to generate relatively unpredictable but smooth trajectory patterns that are consistent across different difficulty levels and participants. The trajectories of are identical within each block but vary across different blocks and participants. The locations (x and y coordinates) of the target and cursor were recorded at sample rate of 60 Hz. Figure 1C shows the target and cursor trajectories of one representative participant in the early and late learning stages at each difficulty level.

#### Task difficulty

In our experiment, the task difficulty is manipulated by varying the cursor radius. In addition to self-reports of subjective workload levels for each task difficulty condition, we propose a measure to quantify difficulty level based on the physical properties of the target and cursor radii. Here, we define 6 relative difficulty levels labeled from L1 to L6, with ID of the *i*^*th*^ difficulty level parameterized as following:

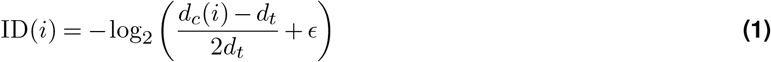

where *d*_*t*_ = 30 mm is the target diameter and *ϵ* = 3 mm is the error term, which are fixed parameters throughout the experiment. *d*_*c*_(*i*) is the cursor diameter of the *i*^*th*^ difficulty level, which is set as *d*_*c*_ ∈ {2.5, 2.0, 1.8, 1.5, 1.3, 1} · *d*_*t*_. According to this definition, a decrease in cursor radius corresponds to an increase in required tracking precision, thereby increasing in task difficulty level. Thus, ID for each task difficulty level is given by ID ∈ {0.23, 0.74, 1.00, 1.51, 2.00, 3.32}. Figure 1B illustrates the relative sizes of the target and cursor for each difficulty level. Visual feedback on tracking performance was given by turning the cursor color from red to green whenever target was within the cursor’s boundary. This condition was met if the euclidean distance (D) between the target and cursor was less than a specified threshold, denoted as 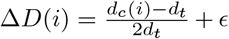 (illustrated as shaded areas in Figure 2A). To distinguish between the two measures of load level evaluation in our experiment, we use the term ‘difficulty level’ to refer to ID, and ‘subjective workload level’ to refer to ISA in the subsequent discussion.

**Figure 1.**
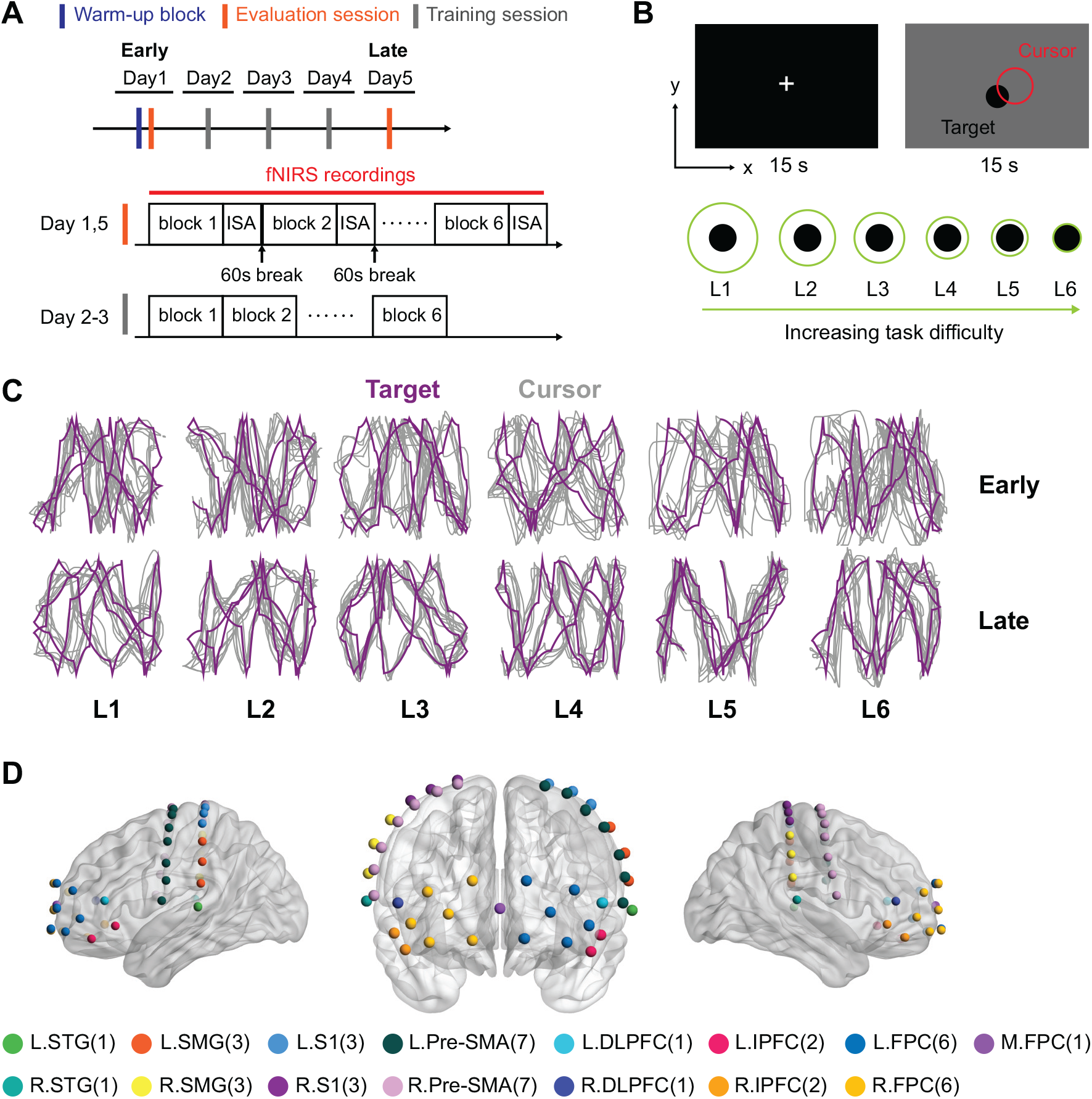
Experimental design. (A) Experimental procedure. Participants visited the lab for consecutive 5 days. After a warm-up block at the beginning of the experiment, participants finished 5 sessions of tasks (1 session/day), including 2 evaluations sessions and 3 training sessions. (B) Continuous target tracking tasks vary in 6 difficulty levels, with increasing task difficulty as the cursor diameter decreased. Performance feedback is given by changing the cursor color from red to green when target was within the cursor’s boundary. The x-axis and y-axis represent the horizontal and vertical directions, respectively. (C) Representative target and cursor trajectories of one participant in the early and late learning stages. Target trajectories are identical within each block but vary across different blocks. (D) Spatial positions of fNIRS channels and ROIs. Dots represent channel positions. 47 channels are clustered into 15 ROIs (labeled in different colors) based on Talairach Daemon labeled Brodmann areas. Numbers in the brackets denote the number of channels in each ROI. Abrreviations: L, left; R, right; M, middle; ISA, instantaneous self-assessment.

### fNIRS data acquisition

Cerebral hemodynamic signals were recorded by a continuous-wave fNIRS device (Nirsmart, Danyang Huichuang Medical Equipment). The system uses two-wavelength (740 nm and 850 nm) light-emitting diode (LED) as light sources with sample rate of 10 Hz. A total of 47 long-separation channels with source-detector distance of 3 cm were created by 23 sources and 15 detectors. The fNIRS cap was symmetrically put on the participant’s head, while the positioning of the fNIRS cap was validated based on a special mark Cz, which should be located at the intersection between sagittal plane and coronal plane on the top of the head.

**Figure 2.**
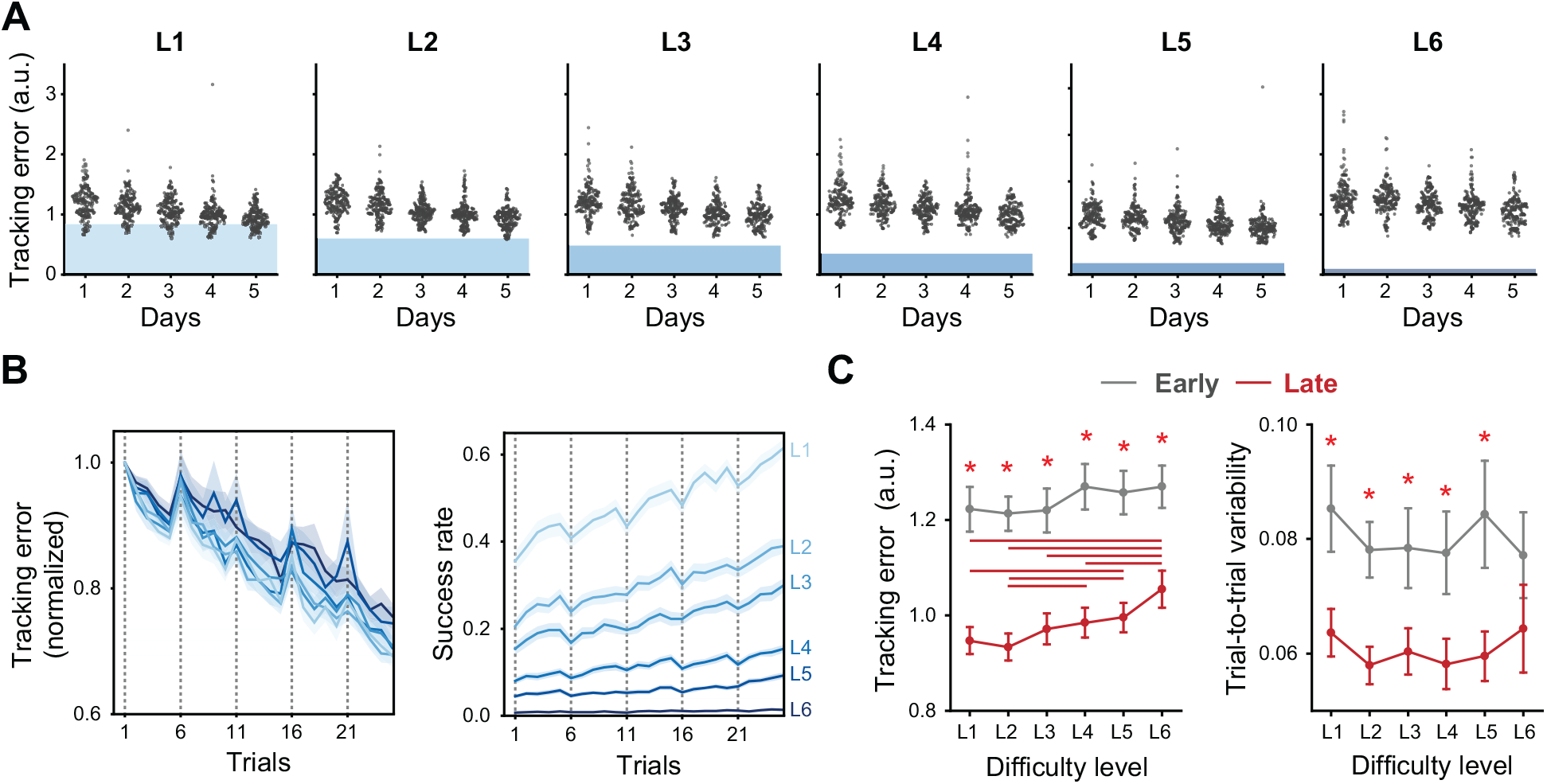
Behavioral results. (A) Tracking errors at each difficulty level throughout the training days. Dots represent individual tracking error within each trial. Green shaded areas indicate the range within which the euclidean distance between the target and the cursor falls. (B) Changes in tracking errors (normalized to the first trial) and success rate across trials at each difficulty level. Shaded areas represent SEM across participants. Vertical dashed lines denote the start of each session. (C) Within-block tracking error and trial-to-trial variability dependent on difficulty level. Red asterisks denote significant difference between late and early learning stages at specific difficulty level. Red horizontal lines denote significant difference (*p <* .05, Bonferroni corrected) between two task difficulty conditions in the late learning stage.

To determine the spatial locations of the channels, we used a Patriot 3D Digitizer (Polhemus, USA) to collect the anatomical locations of all the optodes and 5 reference landmarks (Cz, Nz, Iz, LPA and RPA). The channel locations were then converted to the Montreal Neurological Institute (MNI) coordinates using NIRS-SPM toolbox (Ye et al., 2009). We then clustered the channels into 15 ROIs (7 cortical areas) based on Talairach Daemon labeled Brodmann area (BA) (Lancaster et al., 2000), including frontopolar cortex (FPC, BA-10), dorsolateral prefrontal cortex (DLPFC, BA-46), primary somatosensory cortex (S1, BA-1, 2, 3), pre-motor and supplementary motor area (Pre-SMA, BA-6), inferior prefrontal cortex (IPFC, BA-47), supramarginal gyrus (STG, BA-40), and superior temporal gyrus (SMG, BA-42), as demonstrated in Figure 1D.

### Data analysis

#### Behavioral data analysis

To minimize performance inconsistencies caused by the abrupt initiation of tasks, we omitted the first 1 s data from each trial, focusing our analysis on the remaining 14 s. Behavioral performance is accessed by 3 different measures: tracking error, success rate, and trial-to-trial variability. Tracking error is calculated as the root mean square error (RMSE) between the target and cursor positions within each trial. We also calculated the within-block averaged tracking error as the mean value of the RMSE across 5 trials for each block. Success rate is calculated as the percentage of the time when the target falls within the boundary of the cursor (i.e., the cursor turned green) for each trial. Trial-to-trial variability is calculated as the standard deviation (SD) of RMSE across 5 trial within each block.

To determine how well participants could track the target movements, we evaluated three synchronization measures: coherence, time-lag, and phase-lag. All these evaluations were performed separately for the horizontal and vertical directions. Firstly, we computed the frequency-specific magnitude-squared coherence between the target movement and cursor movement (i.e., participants’ hand response):

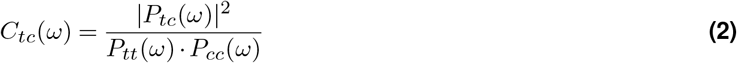

where *p*(*ω*) denotes power spectral density at frequency *ω*. This analysis is conducted using the Matlab function ‘mscohere’. We applied Hamming window with window length of 120 sample points with 50% overlap. This analysis reflects how well cursor movement matches the target movement at different frequency.

Then, we examined whether cursor movement is delayed or advanced to the target movement. Firstly, we estimated the time-lag between target and cursor movement at the maximum of the cross-correlation:

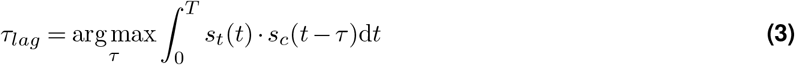

where *s*_*t*_(*t*) and *s*_*c*_(*t*) are the position of the target and cursor, respectively, and *T* is the duration of one trial. Lastly, we analyzed the frequency-specific phase lag based on Fourier transformation:

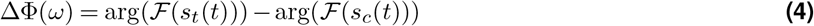

where *ℱ*(·) denotes Fourier transformation, and arg(·) is the argument (phase angle) of a complex number. The value of ΔΦ(*ω*) is unwrapped to correct the phase discontinuity. For both time- and phase-lag estimate, negative values indicate that cursor movement is delayed relative to target movement, and vice versa.

#### fNIRS data analysis

Preprocessing of the fNIRS data are conducted using Homer2 (v2.8) toolbox (Huppert et al., 2009). Due to technical issues (missing or incorrect markers), 11.94% blocks (early: 11.67%, late: 12.22%) are excluded. Here, any exclusion of one trial would cause rejection of the whole block. Poor quality channels are identified based on the absence of *∼*1 Hz cardiac pulsation (Tong et al., 2011), resulting in rejection of 7.27% channels (early: 7.44%, late: 7.09%). Raw data are then converted into optical density, followed by motion artifact correction using spline interpolation method(Scholkmann et al., 2010). Principal component analysis (PAC) is performed to remove global noise (Zhang et al., 2016). Physiological noises and baseline shifts are removed by a three-order Butterworth bandpass filter of 0.01-0.1 Hz. Finally, optical density data are converted into concentration changes of oxyhemoglobin (ΔHbO) and deoxyhemoglobin (ΔHbR) based on modified Beer Lambert’s Law (Baker et al., 2014). Our analyses focus on ΔHbO as it shows superior contrast-to-ration compared with ΔHbR (Strangman et al., 2002).

For individual level analysis, we apply general linear model (GLM) to estimate the task-evoked cortical activation (beta value) using NIRS-KIT toolbox (Hou et al., 2021). Beta values of each participant per each task-level are calculated for subsequent analysis. For ROI-wise analysis, beta values are averaged across channels within each ROI.

### Statistics

We perform statistical analysis using Matlab (version R2020b), JASP (version 0.18.1) and R (version 4.3.1). For all the group-level analysis, outliers are identified as the data points exceed three SD from the mean. To access the influence of difficulty levels and learning stages on behavioral performance, we conduct 2 (learning stages: early, late) × 6 (difficulty levels: L1, L2, L3, L4, L5, L6) two-way repeated measures analysis of variance (rmANOVA) on within-block averaged tracking error and trial-to-trial variability. Sphericity is checked by Mauchly’s test, and no significant violation of the sphericity assumption is found. In the *post-hoc* analysis, multiple comparisons are adjusted using Bonferroni correction.

For group-level comparisons, we use one-sample *t* -test or paired *t* -test, which is indicated in the corresponding results. For group-level cortical activation analysis, we first examine the channel-wise activation by comparing the beta values to zero using one-sample *t* -test. This analysis is conducted separately for each load-level in the early and late stages. Then we use paired *t* -test to estimate the late vs. early differential activation. Multiple comparisons are corrected across 47 channels using Benjamini and Hochberg false discovery rate (FDR) procedure (Benjamini and Hochberg, 1995).

Coefficients of correlation analysis are calculated as either Pearson or Spearman’s rank correlation coefficient, as specified in the corresponding results. For brain-behavior relationship analysis, we estimate the correlation between beta value and within-block averaged tracking error for each ROI separately, and *p* values are adjusted across ROIs using FDR correction.

All data are presented as mean *±* SEM (standard error of the mean), unless otherwise indicated. Statistical significance level is set at 0.05.

### Model comparisons

We apply model-based approach to test the linear versus quadratic effect of load-dependent activation. The linear model assumes linear relationship between cortical activation and load level (i.e., difficulty level, ID, or subjective workload level, ISA), which is written as:

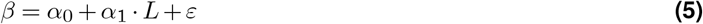

where *β* and *L* denote the cortical activation and load level, respectively. *α*_1_ > 0 indicates cortical activation increases as the load level increases, while *α*_1_ < 0 indicates cortical activation decreases as the load level increases.

For the quadratic model, the relationship between cortical activation and load level is given by introducing an additional quadratic term:

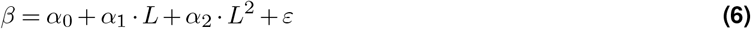

*α*_2_ < 0 indicates inverted U-shaped function, while *α*_2_ > 0 indicates positive U-shaped relationship between cortical activation and load level. Here, we mainly focus on testing whether the load-dependent activation can be better captured by a linear or quadratic model, and and we don’t assume the direction (i.e., positive or negative) of the relationship.

We first fit the models at the group level to examine the load-dependent cortical activation as function of difficulty level (i.e., ID) using linear regression by R function ‘lm’. Then, we test the within-subject load-dependent effect on cortical activation using linear mixed-effects models by R function ‘lmer’. The load-dependent effect is treated as fixed factor, while the subject-specific effect is treated as random factor with random intercepts for each participant. Since load level can be represented by both task difficulty level (i.e., ID) and subjective workload level (i.e., ISA rating), we conduct separate analyses to evaluate how well cortical activation could be predicted by each measure.

We use the Akaike Information Criterion (AIC) to assess the model’s goodness of fit (Burnham and Anderson, 2004). The AIC difference between the two models is given by

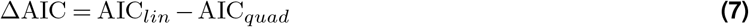

where AIC_*lin*_ and AIC_*quad*_ are the AIC for the linear model and quadratic model, respectively. A model that exhibits a lower AIC value is considered more favorable. As a rule of thumb, an AIC difference greater than 2 indicates substantial evidence favoring the model with the lower AIC, while a difference greater than 10 indicates very strong evidence. Additionally, we use ANOVA test to examine whether one model is significantly better than another.

## Results

### Task difficulty level reflects subjective workload level

To validate the effectiveness of the proposed approach in manipulating load levels in continuous hand movement and how well task difficulty level can reflect subjective load level, we assess the relationship between task difficulty level (i.e., ID) and subjective workload level (i.e., ISA rating). Spearman correlation analysis reveals positive correlations for all the participants in both early and late learning stages, while these correlations increase significantly after training (late vs. early: *t*_(29)_ = 2.436, *p* = .021, paired *t* -test, Figure 3A, B). Moreover, Figure 3C reveals significant positive Spearman’s rank correlation between tracking error and ISA rating in the late learning stage (*rho* = .301, *p* < .001), while the correlations is not significant in the early learning stage (*rho* = .115, *p* = .123). This suggests that better performance (i.e., lower tracking error) is associated with lower subjective workload, and participants knew their performance better after training.

**Figure 3.**
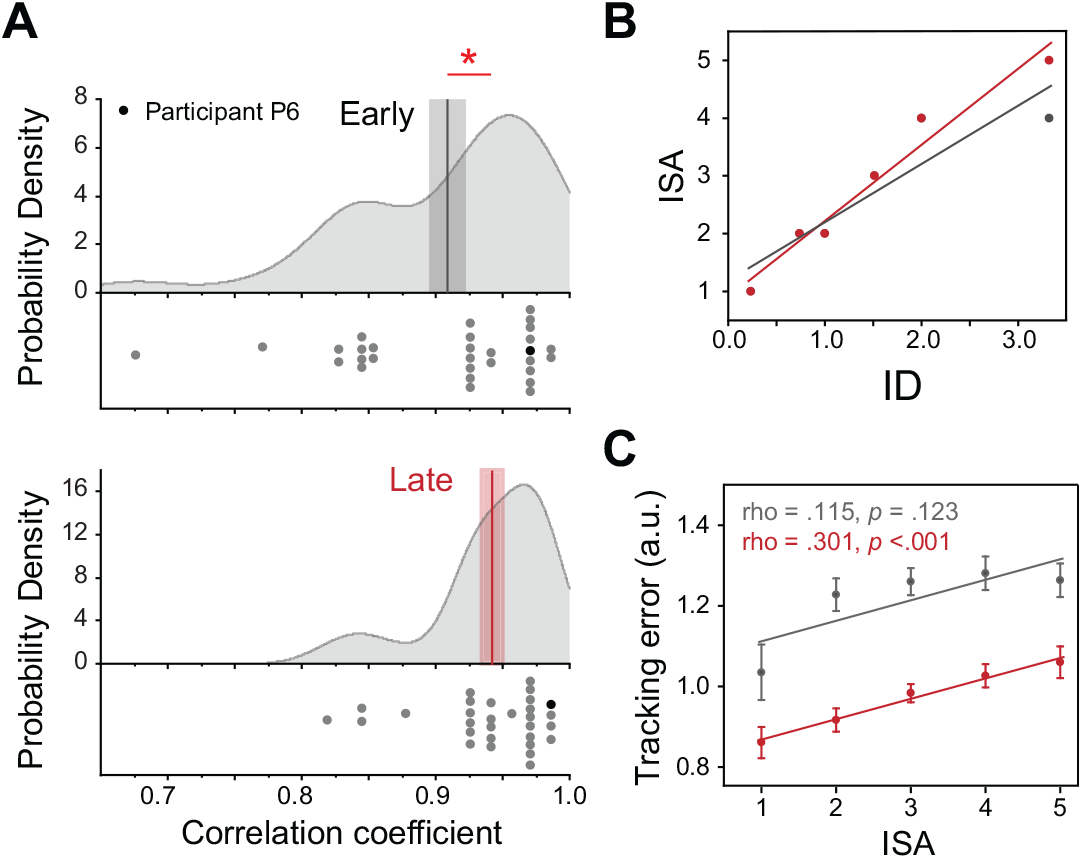
Subjective rating in relation to task difficulty level and behavior performance. (A) Correlation coefficients between ISA and ID at early (upper) and late (bottom) learning stages. Dots indicate individual Spearman correlation coefficients. Red asterisk indicates significant difference in correlation coefficients between late and early learning stage (paired *t* -test, *p <* .05). Shaded areas indicate SEM across participants. (B) Scatter plots of the correlation between ID and ISA rating of one representative participant P6. Lines represent linear regression. (C) Correlation between tracking error and ISA at early and late learning stage. Spearman’s rank correlation. Lines represent linear regressions. Abbreviations: ISA, instantaneous self-assessment; ID, index of difficulty.

### Effective learning after motor training

Behavioral performance is estimated by three main measures: tracking error, success rate, and trial-to-trial variability. Despite the initial decrease in performance at the start of each training session (i.e., overnight forgetting), we observe gradually improving performance (i.e., the decreasing tracking error and increasing success rate) as learning proceeds (Figure 2B), exhibiting a typical pattern of effective learning with deliberate practice (Du et al., 2022).

To evaluate the effect of difficulty level and learning stage on behavioral performance, we perform 2 (learning stage: early, late) × 6 (difficulty level: L1, L2, L3, L4, L5, L6) two-way rmANOVA to test the difference in tracking error and trial-to-trial variability. In terms of tracking error, no significant interaction effect is found between learning stage and difficulty level 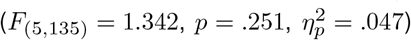. Main effect is found in both learning stage 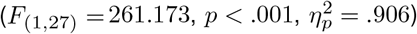 and difficulty level 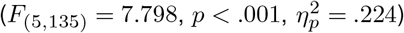. *Post-hoc* analyses reveal significant reduction in tracking errors in the comparisons of late vs. early learning for all difficulty levels (all *p* < .001, Bonferroni corrected), while significant difference across difficulty levels is only observed in the late learning stage (see Figure 2C, left).

For trial-to-trial variability, no significant interaction effect is found between learning stage and difficulty level 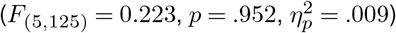. Significant main effect is found in learning stage 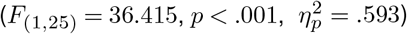, but the main effect in difficulty level is not significant 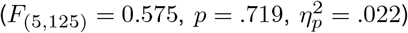. Further, analysis of the effect of learning stage indicated significantly reduced trial-to-trial variability in the late vs. early stage of learning for difficulty levels from L1 to L5 (all *p* < .05, Bonferroni corrected), but no significant difference is found in effect of difficulty level for both early and late learning stages (see Figure 2, right).

**Figure 4.**
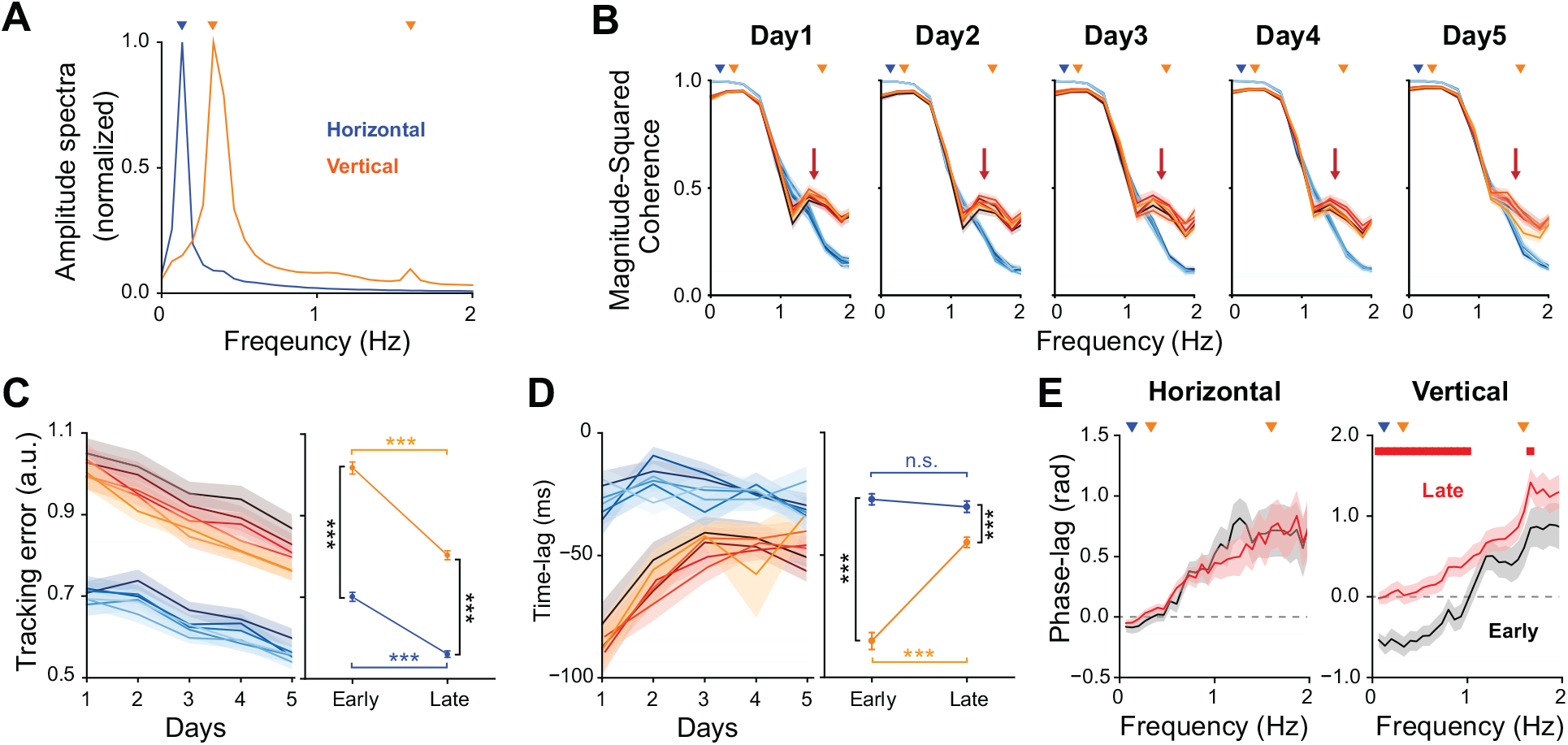
Directional-specific behavioral results. (A) Frequency-domain amplitude spectra, averaged across conditions and participants, for target movements in the horizontal (blue) and vertical (orange) directions, respectively. Downwards triangles denote amplitude peaks. (B) Frequency-specific coherence between the target and the hand (i.e., the cursor) movements are separately analyzed for horizontal (blue) and vertical (orange) direction. Red arrows indicate a diminishing peak at high frequency as training progresses. Downwards triangles denote amplitude peaks in the the amplitude spectra of target movements. (C) Tracking errors in the horizontal (blue) and vertical (orange) directions respectively (left). Darker colors indicate higher levels of task load. Tracking error is collapsed across different load levels (right). (D) Time-lag between the target and hand movements in the horizontal (blue) and vertical (orange) directions respectively (left). Time-lag is collapsed across different load levels (right). ^*\*\*\**^*p <* .001, n.s., not significant. Two-way rmANOVA, *post-hoc* analysis using paired *t* -test with Bonferroni corrected. (E) Phase-lag between the target and hand movements in the horizontal (left) and vertical (right) directions, collapsed across all load levels. Red dots denote significant difference between early and late learning stages (*p <* .05, paired *t* -test). Negative values of time- and phase-lag indicate the hand movements are delayed relative to the target movements, and vice versa. Error bars and shaded areas indicate SEM.

### Characteristics of target-hand synchronization are directional-specific and load-independent

To understand how well participants could track the target movements, we assess the synchronization between the target and hand trajectories. Figure 4A shows the amplitude spectra of the target movements, averaged across conditions and participants, for the horizontal and vertical directions, respectively. In the low-frequency range (0-1 Hz), there are peaks at 0.13 Hz for the horizontal direction and 0.33 Hz for the vertical direction. Additionally, in the high-frequency range (1-2 Hz), there is a peak at 1.60 Hz in the vertical direction. As demonstrated in Figure 4B, the magnitude-squared coherence between the target and hand movements reveals high coherence at low-frequency and low coherence at high-frequency, suggesting that the participants can follow the overall trends of the target movements (slow and smooth movements) but are unable to track the rapid changes in the target movement. Meanwhile, coherence in the high-frequency range is higher in the vertical direction compared to the horizontal direction, with peaks around 1.60 Hz that match the peak amplitude spectra of the horizontal target movements. As learning progresses, these high-frequency coherence peaks in the vertical direction diminish (indicated by red arrows in Figure 4B), even though the overall tracking accuracy improves over time. This suggests that as participants become more familiar with the task, they may adopt more efficient tracking strategies by focusing more on the overall trends of target movements and neglecting the rapid motion. Notably, the directional-specific patterns of target-hand coherence are consistent across different difficulty levels.

Participants also exhibit directional asymmetry in behavioral performance, consistently showing higher tracking error in the vertical direction (Figure 4C). Specifically, we find significant interaction effect between direction and learning stage 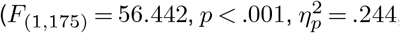, two-way rmANOVA), and main effects of direction 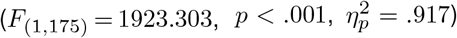 and learning stage 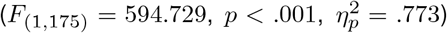. We then examine whether the hand movements are delayed or advanced relative to the target movement. For both horizontal and vertical directions, participants’ hand movements are delayed relative to the target. However, the patterns of lag-time differ between directions across different training sessions: the lag-time remains consistent in the horizontal direction, while it gradually increases (i.e., less delay) in the vertical direction (Figure 4D). Two-way rmANOVA analysis reveals significant interaction effect between direction and learning stage 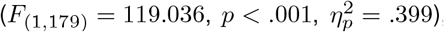, and main effects of direction 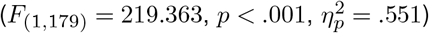 and learning stage 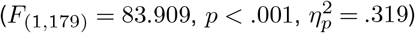.

Further, we analyze the frequency-specific characteristic of phase-lag, focusing on the differences between the early and late learning stages. Overall, participants show an increasing trend of phase-lags as frequency increases, reflecting an tendency towards predictive control strategies at higher frequencies. In the horizontal direction, the phase-lags are positive across the the whole frequency range, and this pattern remains consistent in both early and late learning stages (Figure 4E, left). By contrast, in the vertical direction, the overall phase-lags increase with effective learning. More specifically, significant increase (from negative to positive) in phase-lag after training is observed in the low-frequency range, showing clear evidence of a shift from corrective (i.e., feedback) towards predictive (i.e., feedforward) control strategies. In the high-frequency range, phase-lags are positive in both the early and late learning stages, with a significant difference found only at 1.666 Hz, the frequency around where the target’s vertical movement amplitude spectrum peaks (Figure 4E, right).

**Figure 5.**
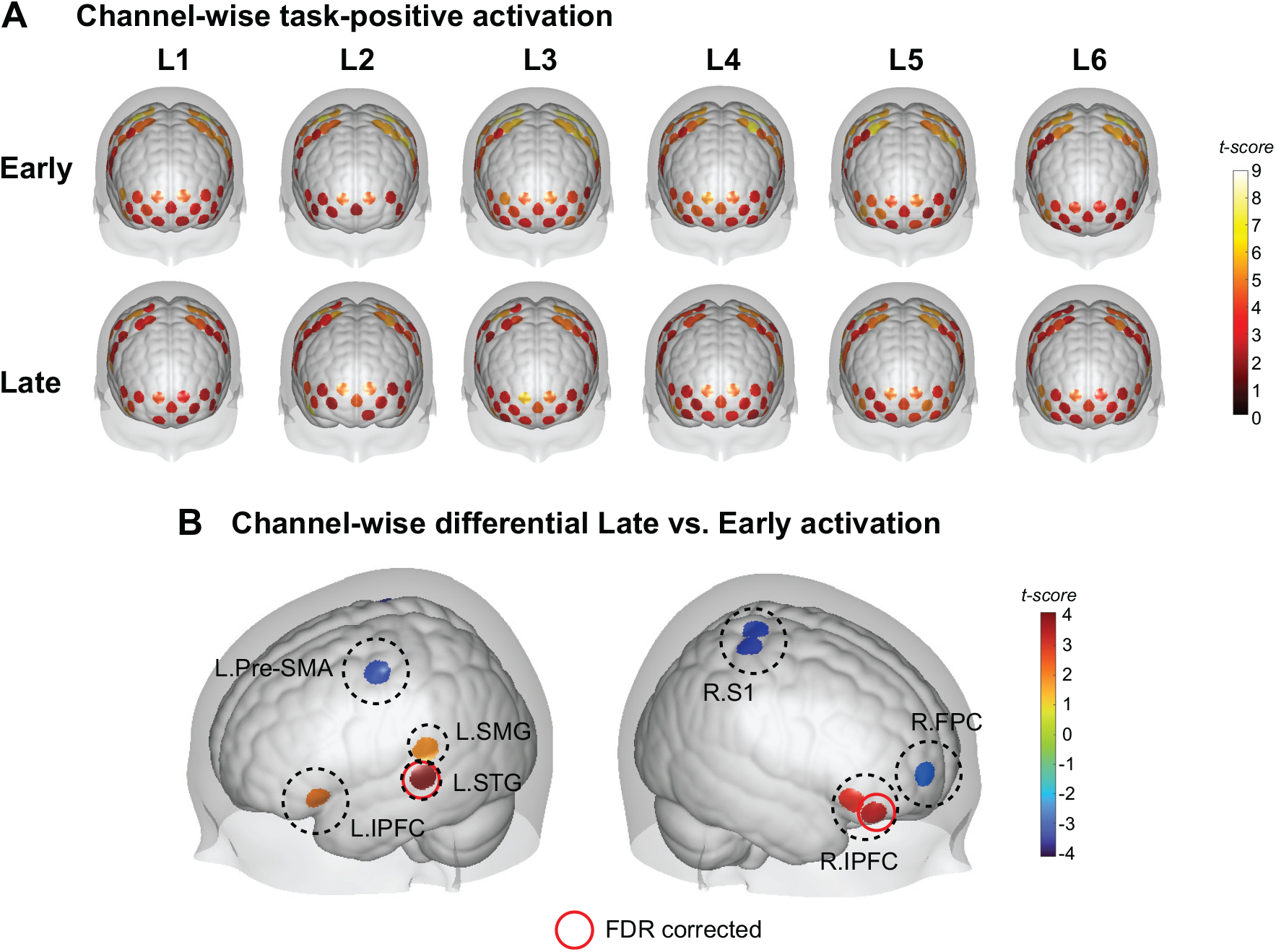
Channel-wise cortical activation results. (A) Task-related activation per load level at early and late learning stage. Regions scaled by red colors represent beta values significantly greater than 0. One-sample *t*-test, *q*_(*FDR*)_ *<* .05. (B) Differential activation of late vs. early learning stages. Beta values are collapsed across load levels for each learning stage. Regions scaled by red colors indicate significant increased activation during late learning stage compared with early learning stage, while blue colors indicate significant decreased activation during late learning stage. Paired *t*-test, *p <* .05, uncorrected; red circles denote *q*_(*FDR*)_ *<* .05.

### Dissociation of activity in the fronto-pariatal network with effective motor skill learning

To estimate the channel-wise cortical activation, we perform exploratory analysis for all channels and conditions. We compare the beta values to 0 using one-sample *t* -test for each channel, which is conducted for each load level and learning stage separately. Results show widespread task-positive (beta value greater than 0) activation across the fronto-parietal cortical areas in all conditions (Figure 5A). We then collapsed the beta values across load levels, mainly focusing on the differential activation of late vs. early learning stages. Significant increased activation is found in left STG and right IPFC (paired *t* -test, *q*_*FDR*_< .05), and left IPFC and left SMG (paired *t* -test, *p* < .05, uncorrected) after training. Additionally, decreased activation is found in left Pre-SMA, right S1 and right FPC (paired *t* -test, *p* < .05, uncorrected) after training (see Figure 5B).

**Figure 6.**
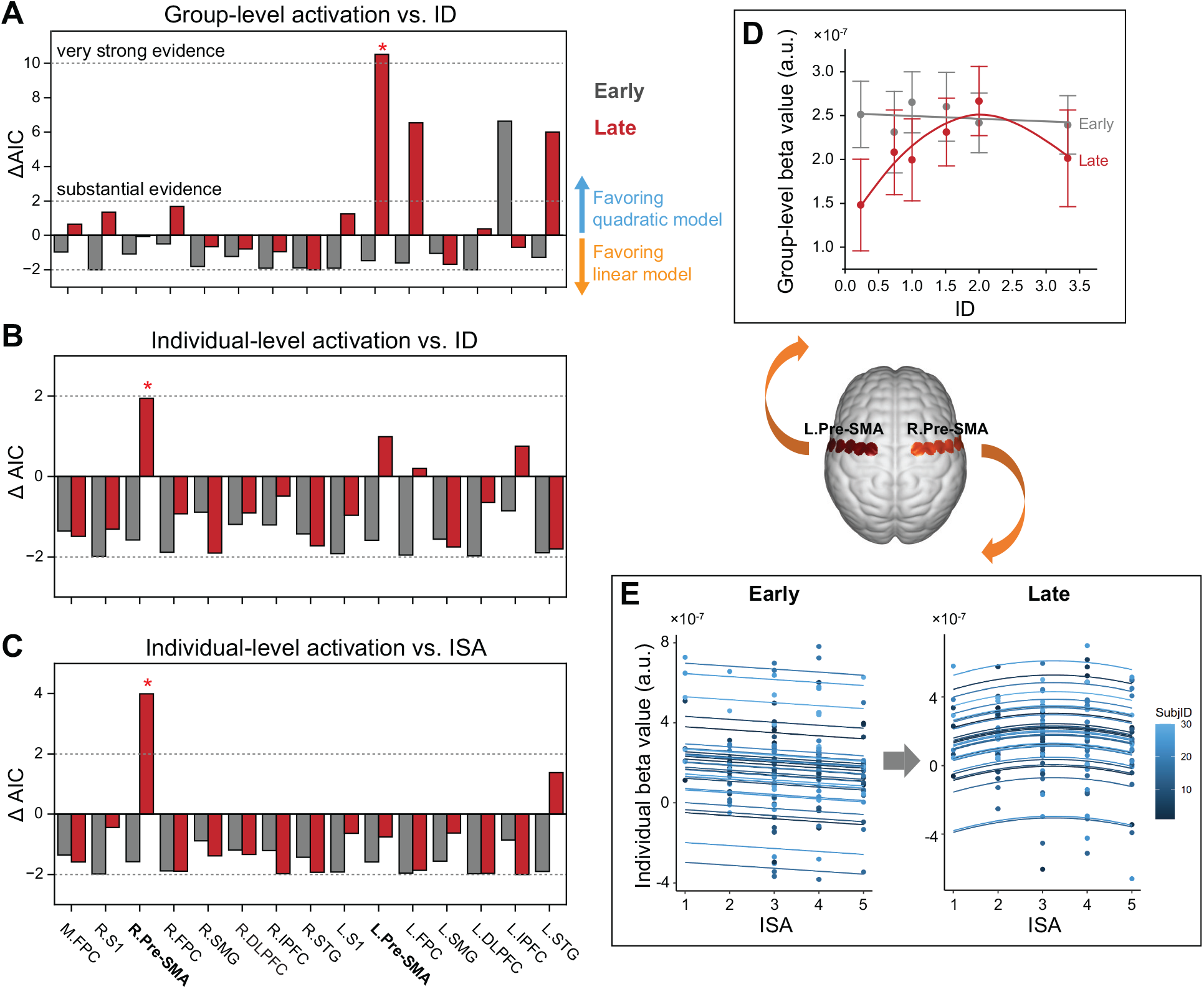
Learning-induced changes in load-dependent cortical activation. Linear vs. quadratic model comparisons are performed in three measures: (A) Group-level activation predicted by ID, (B) individual-level activation predicted by ID, and (C) individual-level activation predicted by ISA. Positive ΔAIC favors the quadratic model over the linear model. AID difference exceeding 2 is typically regarded as the threshold for substantial evidence, while a difference greater than 10 is considered indicative of very strong evidence in favor of one model over another. Red asterisk indicates significant difference between linear and quadratic model (ANOVA, *p <* .05). (D) Group-level cortical activation as a function of ID in left Pre-SMA in the early stage (predicted by linear model) and late stage (predicted by quadratic model). (E) Individual-level cortical activaiton as function of ISA in right Pre-SMA in the early stage (predicted by linear model) and late stage (predicted by quadratic model). Solid lines represent regression results fitted by either linear or quadratic models. Error bars indicate SEM. Abbreviations: ISA, instantaneous self-assessment; ID, index of difficulty.

### Load-dependent cortical activation

To test whether linear or quadratic model can better capture the relationship between cortical activation and load level, we conduct model comparison for both group and individual level effects. Overall, no substantial evidence supporting the linear model is found regardless of ROIs and learning stages, with all negative ΔAIC not exceeding -2 (Figure 6A-C). For group-level effect, very strong evidence favoring quadratic model is found in left Pre-SMA in the late learning stage (ΔAIC = 10.512, *F*_(1,3)_ = 21.144, *p* = .019). Regarding the individual-level effect, we fit individual-level activation as function of either ID or ISA, with within-subject effect as random factor. In the late learning stage, the relationship between cortical activation and load level is better captured by the quadratic model in right Pre-SMA for both ID (ΔAIC = 1.947, χ^2^(1) = 3.947, *p* = 0.047) and ISA (ΔAIC = 3.992, χ^2^(1) = 5.992, *p* = .014). Note that the load-dependent pattern is better captured by the linear model in the early stage, although with weak evidence (AIC difference less than 2). Our results reveal significant shifts towards quadratic relationship between cortical activation and load level in Pre-SMA after training.

We then analyze the specific load-dependent patterns in Pre-SMA, focusing on the left Pre-SMA at group-level and right Pre-SMA at individual-level in different learning stages. For both group and individual level analyses, we observe decreasing cortical activation as load level decreases in the early stage. By contrast, in the late stage, the load-dependent activation exhibit inverted-U shape pattern(Figure 6D, E). These findings indicate that effective motor skill leaning may elicit alteration in the load-dependent activation patterns, leading to the emergence of inverted-U shape relationship between cortical activation and load level. This phenomenon is particularly prominent in Pre-SMA, suggesting the development of the neural efficiency to exhibit optimal activation pattern.

### Brain-behavior relationship

By estimating the Pearson correlation coefficients between beta values and tracking errors, we find negative correlation between cortical activation and tracking error is found in all ROIs and in both early and late learning stages (see Figure 7A). This indicates that better performance (i.e., lower tracking error) is associated with higher cortical activation, suggesting high neural capacity for higher performers – the ability for high performers to intentionally sustain maximal activation even in challenging tasks (Barulli and Stern, 2013; Heinzel et al., 2014) Specifically, in the early stage of learning, significant negative correlation is observed in the right Pre-SMA, right IPFC, left STG, and bilateral DLPFC (all *q*_*FDR*_< .05). As learning progresses to the late stage, these correlations persist, and additional significant correlations emerges in other areas, including the right S1, right STG, bilateral FPC and bilateral SMG (all *q*_*FDR*_< .05). This expansion in the areas that exhibit strong brain-behavior correlation implies an evolving and broadening neural network engagement, highlighting the brain’s adaptive changes in supporting skill acquisition.

**Figure 7.**
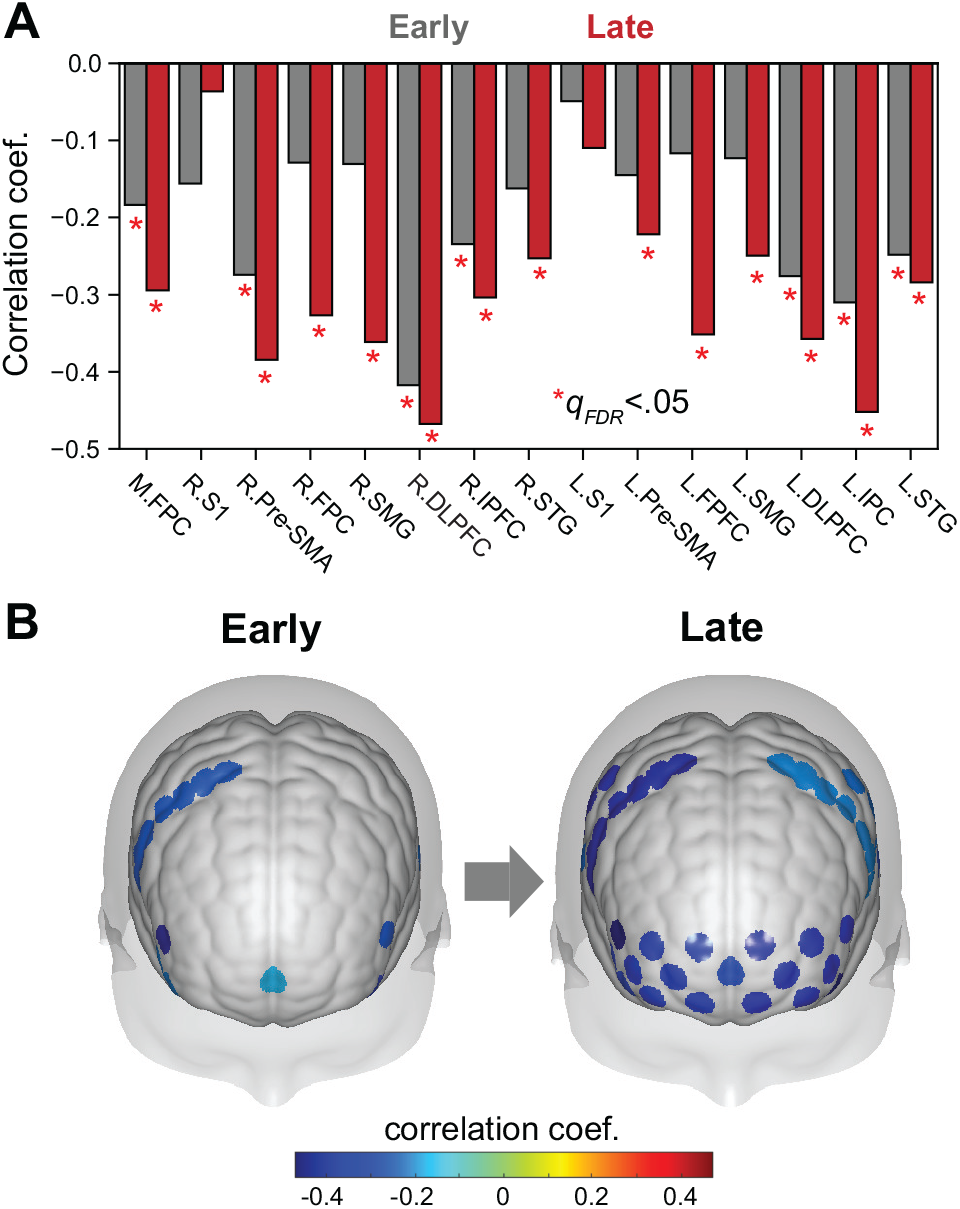
Individual difference of brain-behavior relationship. (A) Correlations coefficients between cortical activation and within-block averaged tracking error for each ROI in the early and late learning stages, respectively. Negative correlations indicate that high performance (i.e., low tracking error) individuals exhibit greater cortical activation. Correlations coefficients are estimated as partial Pearson correlation with age as covariates. Red asterisks denote significant correlations, with FDR corrected across ROIs. (B) More brain areas exhibit significant brain-behavior relationship in the late learning stage.

## Discussion

In this study, we explore the effect of load level on motor learning in continuous hand movements by a longitudinal fNIRS study. Here, the load level is represented by a quantitative measure of task difficulty level in continuous target tracking tasks. This measure is validated to positively correlate with psychometric measurement of subjective workload level. Crucially, our results demonstrate load-dependent and load-independent accounts associated with effective motor skill learning. Specifically, we show that the control strategy shifts from feedback-dominant to feedforward-dominant as learning progressed, which is load-independent (i.e., consistent across different load levels). Meanwhile, the load-dependent patterns of task-relevant neural response are region-specific and learning-stage-specific, while effective motor skill learning could elicit shifts towards inverted-U relationship between cortical activation and load level, particularly evident in the pre-motor and supplementary areas.

### Enhanced self-monitoring after training

The well-established Fitts’s ID has been consistently shown to be an effective measure for performance predicting and measuring in discrete target pointing tasks (Langolf et al., 1976; MacKenzie, 2018). In this study, we adapt Fitts’s ID to the scenario of continuous target tracking tasks, where the proposed ID increases as the cursor size decreases. Crucially, we find that this adjusted ID significantly correlates to subjective mental workload (Figure 3A, B), suggesting that by varying the cursor size can effectively manipulates subjective load levels. Moreover, enhanced correlation is found in the late learning stage, possibly because participants became more familiar with the various levels of task difficulty as training progressed, so they were able to utilize the full range of the rating scale more efficiently. We also observe significant correlation between tracking error and ISA in the late learning stage (Figure 3C). These results suggest that participants were able to assess their performance more accurately with practice, reflecting enhanced self-monitoring after training. This is possibly associated with the strengthened brain-behavior relationship after training (Figure 7), while the brain encodes information more efficiently in the late learning stage, allowing participants to have better cognitive control and attention to their internal states.

### Transition from feedback-dominant to feedforward-dominant control as learning progressed

Previous studies has demonstrated that the integration of feedback (i.e., corrective) and feedforward (i.e., predictive) control strategies play crucial roles in achieving accurate motor movements in a visually-guided target tracking task (Wolpert et al., 1995; Scott, 2016; Kawato, 1999; McNamee and Wolpert, 2019; Wagner and Smith, 2008; Maeda et al., 2018, 2020). Here, we provide clear evidence of the transition from feedback-to feedforward-dominant control during longitudinal motor learning. The patterns of tracking strategies at different training sessions are load-independent (i.e., consistent across different load levels) and direction-specific (i.e., inconsistent between the horizontal and vertical directions).

As training progressed, participants’ hand movements in the vertical direction became less correlated with the target movements at high-frequency range (Figure 4B). Moreover, Figure 4E shows transition from negative phase-lag (i.e., hand movements are delayed relative to the target) to positive phase-lag (i.e., hand movements are advanced relative to the target). These results indicate that participants may adopt a less corrective but more predictive control strategy in the late learning stage. Accordingly, we observe decreased activity in the S1 and Pre-SMA areas after training (Figure 5B), suggesting less engagement of feedback processing and active motor planning, but more reliance on predictive control when motor execution becomes more automatic and efficient (Poldrack et al., 2005; Floyer-Lea and Matthews, 2004). Indeed, anecdotal evidence from participants suggests the potential use of a feedforward control strategy in the late stage of learning: Several participants reported that they’ve noticed that the trajectories remained consistent within each block. Therefore, they attempted to anticipate the target’s movement and employed an interceptive strategy to capture the target.

The feedback and feedforward interaction could also attribute to the balance between exploration and exploitation in motor learning and adaptation (Dhawale et al., 2017; Krakauer et al., 2019). Our results show decreased trial-to-trial variability after training (Figure 2C), suggesting a transition from exploration-to exploitation-dominant strategy with training. In the early stage, participants exhibited more unsuccessful movement and they would explore possible better movements that could catch the target and hence achieve higher rewards. In this stage, they were more likely to perform corrective movements based on sensory feedback. As training progressed, the motor system can exploit the success in precise tracking movements and try to reproduce similar movements as much as possible, thereby reducing variability. This transition in learning strategy is associated with decreased activity in right FPC (Figure 5B), an area known be causally involved in the exploration-exploitation trade-off (Raja Beharelle et al., 2015; Zajkowski et al., 2017).

It’s noteworthy that the patterns of feedback-feedforward transition are only observed in the vertical direction. This is possibly due to the directional asymmetry in the smooth pursuit eye movements, as more accurate tracking performance is found in the horizontal direction for both the eyes and hand (Ke et al., 2013; Danion et al., 2021). In line with previous findings, we observe more accurate tracking performance in the horizontal direction throughout the training periods (Figure 4C). Due to the inherent advantages in the horizontal movements, participants consistently exhibited higher tracking accuracy. In contrast, participants initially showed lower tracking accuracy in the vertical direction. With repeated practice, their proficiency in this direction improved, exhibiting increased accuracy and reduced delays (or even advance) in the late stage. Despite these improvements, vertical performance remained inferior to horizontal performance due to the underlying structural and functional constraints. However, we cannot rule out the influence of different frequency components in the horizontal and vertical directions, and further studies would be needed to address these direction-specific effects. Nonetheless, our findings demonstrate clear evidence of the learning-induced transition from feedback-to feedforward-dominant control, which is consistent across different load levels.

### Learning-induced inverted-U shape pattern of load-dependent activation

We observe widespread bilateral task-positive activation in the frontoparietal areas for all experiment conditions (Figure 5A). Although it is commonly believed that unimanual motor execution is predominantly controlled by the contralateral hemisphere, some evidence suggests that bilateral activation occurs during unimanual hand movements (Chettouf et al., 2020, for review). Previous evidence has identified ipsilateral activation of mirror neurons in the Pre-SMA, S1, and the IPFC (Molenberghs et al., 2009). The broad activation in the frontoparietal network support information processing of sensory feedback and integration, and hence facilitate precise motor control and visual-motor coordination. Both the increase and decrease activity after training suggest learning-induced reorganization in the task-relevant brain networks (Poldrack, 2000; Dayan and Cohen, 2011; Sigman et al., 2005).

How does brain activation in these task-relevant network vary across different load levels, and does learning alter the pattern of load-dependent activation? Previous work has demonstrated the relationship between neural activity and task difficulty as increasing, decreasing or inverted U-shape function (Barch et al., 1997; Gevins, 1997; Charroud et al., 2015; Engineer et al., 2012; Meidenbauer et al., 2021). Additionally, dissociation in the decrease/increase activation with increasing task load has been observed across different brain network in cognitive tasks (Gevins, 1997; Charroud et al., 2015). In the current study, our results provide clear evidence that the load-dependent patterns of brain activation are brain region-specific and may vary in different learning stages of motor learning. Specifically, effective motor skill learning could lead to alteration in the load-dependent patterns of cortical activation. More importantly, the load-dependent neural activity is unlikely to be driven by a linear model (i.e., a simple decreasing or increasing effect). Rather, learning may facilitate the detectability of the inverted-U pattern, as we identify robust evidence of the shift towards inverted-U shaped pattern of the load-dependent cortical activation in the late learning stage, particularly evident in Pre-SMA.

### Real-world implications

The current study has important implications for real-world applications. Firstly, we propose a simple but effective approach for quantifying subjective workload in continuous motor tasks. This provides a scenario closely mirroring real-world situations, holding important potentials for deepening understanding motor skill acquisition mechanisms. Previous neurofeedback studies has suggested that by adapting task difficulty to an optimal load level can facilitate behavioral outcomes (Faller et al., 2019; Zheng et al., 2022). To this end, our quantitative description of subjective workload could enable precise manipulation in such load-adaptive motor training system. Interestingly, a prior study indicated that building a neurofeedback system at appropriate mental workload level could significantly enhance the system’s classifiability (Ke et al., 2016). This poses a challenge for developing training protocols suitable for real-world applications, where the accurate and robust detection of mental workload is crucial. Here, our findings provide evidence of the learning-induced alteration in the load-dependent cortical activation and brain-behavior relationship patterns, revealing complex brain dynamics as motor skill learning progresses. Therefore, it would be an intriguing future research to determine whether and how to leverage these load-dependent motor learning mechanisms to enhance the effectiveness of neurofeedback training and rehabilitation programs.

