## Supplementary figures and images for "Load-dependent and load-independent effects on longitudinal motor training in human continuous hand movements"

### Extended Figure4-1

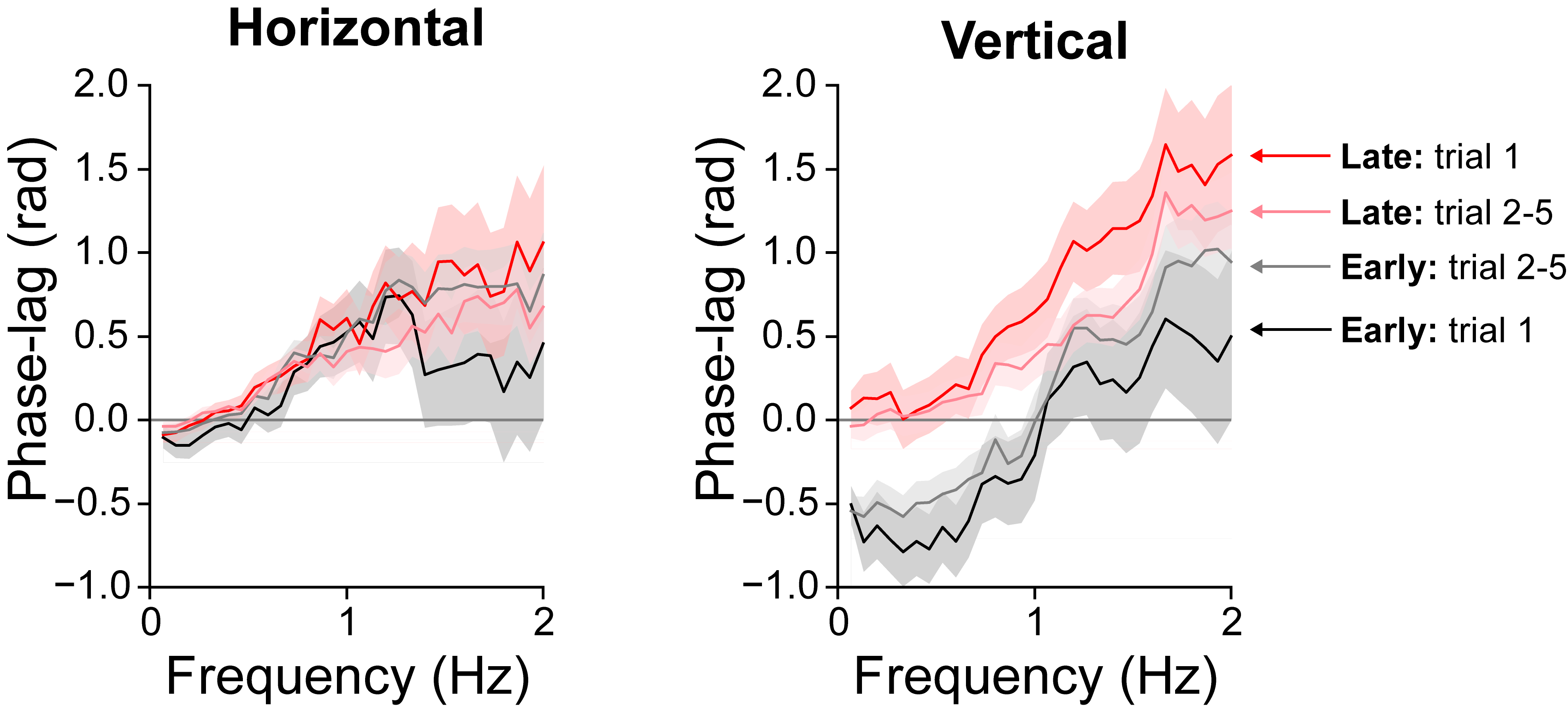
